# PLncFire enables genome wide identification and annotation of plant long noncoding RNAs from RNA sequencing data

**DOI:** 10.1101/2025.10.13.682253

**Authors:** Surbhi Mistry, Shivani Saxena, Ahsan Z Rizvi

## Abstract

**Background:** Long non-coding RNAs (lncRNAs) play important regulatory roles in plant growth, development, and stress responses. However, their genome-wide identification remains challenging due to low sequence conservation, incomplete reference annotations, and variability across species. Existing workflows often lack standardization and reproducibility, limiting large-scale comparative studies. To address these challenges, we developed PLncFire, a modular computational pipeline designed for automated and reproducible identification and annotation of plant lncRNAs using RNA-seq data.

**Methods:** PLncFire processes standard RNA-seq datasets through a structured workflow comprising quality control, read alignment, transcript assembly, and transcript filtering. A consensus-based coding potential assessment strategy was implemented using CPC2, PlantLncPipe, and FEELnc to improve prediction reliability. Transcripts were filtered based on length, exon structure, and coding probability thresholds to generate high-confidence lncRNA candidates. Identified lncRNAs were further classified as known or novel by comparison with reference annotations. The pipeline also incorporates differential expression analysis to support functional prioritization. Workflow modularity ensures scalability across plant species and enables reproducible execution in diverse computational environments.

**Results:** Application of PLncFire to plant RNA-seq datasets enabled systematic identification of high-confidence lncRNA candidates, including both previously annotated and novel transcripts. The consensus coding-potential framework reduced false-positive predictions compared to single-tool approaches. Integration of differential expression analysis facilitated prioritization of lncRNAs associated with specific developmental stages or stress conditions. The modular design demonstrated compatibility across datasets from different plant species, supporting cross-species adaptability and comparative analysis.

**Conclusions:** PLncFire provides a standardized and reproducible framework for genome-wide lncRNA discovery in plants. By integrating multi-tool consensus coding assessment with transcript assembly and expression analysis, the pipeline enhances prediction confidence and scalability. This platform supports large-scale functional genomics studies and facilitates systematic exploration of plant lncRNA landscapes. The source code is available at https://github.com/ahsan-rizvi/PLncFire.git.

**Trial registration:** Not applicable.

## 1 Introduction

Long non-coding RNAs (lncRNAs) are transcripts longer than 200 nucleotides that do not encode proteins, yet play vital roles in regulating gene expression (Zhang et al., 2019; Budak et al., 2020; PSVV et al., 2025). In plants, lncRNAs have been associated with key biological processes, including flowering time control, tissue differentiation, stress adaptation, and epigenetic regulation (Jha et al., 2020). The growing catalogue of plant lncRNAs uncovered through high-throughput sequencing highlights their abundance and functional diversity but also underscores a challenge: unlike protein-coding genes, lncRNAs show poor sequence conservation, variable structures, and highly context-specific expression patterns (Shukla et al., 2021; Zhang et al., 2023). These characteristics make their genome-wide discovery and annotation far from straightforward (Chi et al., 2019; Karmakar et al., 2019). While numerous computational pipelines for lncRNA discovery exist, most have been designed with animal genomes in mind, lack plant-specific optimizations, or depend on a single prediction algorithm, leading to incomplete or error-prone results (Ramakrishnaiah et al., 2020). Furthermore, many approaches stop at identifying putative lncRNAs without integrating downstream classification into known and novel categories or linking them to functional prioritization steps such as differential expression analysis (Mistry et al., 2026). The absence of a standardized, modular, and reproducible workflow tailored to plant systems has limited cross-species comparisons and slowed progress in functional genomics.

For instance, PlantLncPipe (Tian et al., 2024) offered one of the first end-to-end workflows for plants, but its reliance on a single coding-potential predictor limited both its accuracy and generalizability. FEELnc (Wucher et al., 2017), widely adopted across species, was originally optimized for animal genomes and often misclassifies plant transcripts due to their repetitive structures and incomplete annotations. CNIT (Guo et al., 2019) applied alignment-free features to improve speed but struggled with interpretability and cross-species reproducibility. PLncPRO (Singh et al., 2017) introduced a random forest framework, yet its predictions varied widely depending on input annotation quality. More recently, deep learning–based models such as DeepPlnc (Ritu et al., 2021) demonstrated improved sensitivity but at the cost of high false-positive rates when applied to complex plant transcriptomes. The latest entrant, PlantLncBoost (Tian et al., 2025), was marketed as a breakthrough; however, an initial version misclassified chromosome identifiers as lncRNAs, and the tool continues to suffer from severe overprediction and an unstable precision-recall balance despite updates.

Collectively, these advances highlight the growing interest in computational lncRNA discovery, but they also underscore a persistent gap: the absence of a modular, statistically robust, and biologically interpretable pipeline tailored specifically to the complexities of plant genomes. To address this, we introduce PLncFire, a comprehensive and specialized computational pipeline designed for the robust identification and annotation of long non-coding RNAs (lncRNAs) in plant transcriptomes. To overcome the critical challenges of high false-positive rates and poor cross-species conservation, PLncFire integrates a multifaceted strategy. It begins with stringent pre-processing of sequencing data to ensure a high-quality foundation, followed by a robust consensus prediction phase that leverages the complementary strengths of three distinct tools: CPC2 (Kang et al., 2017), PlantLncPipe (Tian et al., 2024), and FEELnc (Wucher et al., 2017). This ensemble method significantly enhances prediction reliability by minimizing the biases and limitations inherent to any single algorithm. Subsequently, each putative lncRNA is annotated through meticulous searches against curated databases, classifying them into known or novel categories.

Architecturally, this pipeline is designed to be modular and scalable, ensuring it can be readily adapted to diverse plant species with varying genomic complexities and to data from different sequencing platforms. Most importantly, PLncFire moves beyond mere identification by seamlessly integrating downstream functional prioritization, such as differential expression analysis, directly into its workflow. Consequently, PLncFire does not merely generate a catalogue of candidates; it provides a powerful, standardized framework for the functional exploration of lncRNAs, thereby accelerating research into their roles in plant development, stress response, and evolutionary biology.

What distinguishes PLncFire is its consensus-based design and its ability to provide consistent performance across the evaluated datasets. Unlike existing pipelines that either miss a substantial fraction of reference lncRNAs or inflate predictions with false positives, PLncFire achieves a critical balance, consistently recovering annotated transcripts while maintaining high precision, recall, and F1 scores. Similar performance trends were observed across the evaluated datasets, suggesting that the workflow can be applied reproducibly under the tested conditions. By combining multiple complementary prediction tools within a consensus framework, PLncFire aims to improve confidence in lncRNA identification by reducing dependence on any single prediction method.The pipeline’s scope extends well beyond initial discovery: its modular framework supports comparative analyses, cross-species studies, and integration with functional genomics data, making it a versatile tool for advancing both basic and applied plant science.

## 2 Materials and Methods

The PLncFire pipeline was developed as a modular, transcriptome-based framework for the systematic identification and annotation of plant lncRNAs. The workflow begins with rigorous data pre-processing to ensure that only high-quality reads enter downstream analyses. Raw RNA-seq reads were first subjected to quality assessment with FastQC (Brown et al., 2017) to evaluate sequencing integrity, adapter contamination, and base composition. Low-quality bases and adapters were removed using Trimmomatic-0.39 (Bolger et al., 2014) with default parameters. Genomes were indexed with HISAT2-build (Guo et al., 2022), and quality-trimmed RNA-seq reads were aligned to the reference genome. The resulting alignment files were sorted and indexed with SAMtools (Li et al., 2009; Danecek et al., 2021),to provide a stable and reproducible basis for transcript reconstruction. Transcriptome assembly was performed using StringTie (Shumate et al., 2022) in reference-guided mode, enabling the recovery of both annotated and potentially novel transcripts. Assemblies from multiple samples were merged to generate a comprehensive, non-redundant transcript catalogue. To refine this set, transcripts shorter than 200 nucleotides or those containing fewer than two exons were removed, and overlaps with annotated protein-coding genes were excluded using Gffcompare (Pertea and Pertea, 2020), thereby enriching the dataset for putative non-coding candidates. To achieve reliable lncRNA identification, the workflow combines several prediction tools, mitigating the elevated false-positive rates (Tian et al., 2025) associated with individual methods.

PLncFire scalable pipeline implements a consensus-based strategy for coding potential assessment. Candidate transcripts were evaluated using three complementary tools: CPC2 (Kang et al., 2017), which relies on sequence-intrinsic features; PlantLncPipe (Tian et al., 2024), which applies plant-optimised classifiers; and FEELnc (Wucher et al., 2017), which incorporates machine learning and comparative annotation. Only transcripts classified as non-coding by at least two of these three tools were retained, substantially reducing false positives while capturing genuine lncRNAs. This consensus approach represents a central innovation of PLncFire, setting it apart from existing plant-specific pipelines. Following identification, the retained candidates were annotated by homology searches against curated plant lncRNA databases, including RNAcentral (Sweeney et al., 2019), CANTATAdb (Szcześniak and Wanowska, 2024), and PLncDB (Jin et al., 2021), using BLASTn (Camacho et al., 2009). Transcripts with significant matches were assigned as known lncRNAs, while unmatched candidates were considered novel, thus establishing a reproducible framework for distinguishing conserved regulators from lineage- or condition-specific transcripts.

PLncFire scalable schematic is summarised in Figure 1, follows a modular design comprising three stages: pre-processing, identification, and annotation with prioritisation. Each stage is fully containerised, ensuring reproducibility across platforms and compatibility with both local workstations and high-performance computing environments. In addition, the modularity of PLncFire allows substitution of individual tools; for instance, HISAT2 (Guo et al., 2022) can be replaced by STAR (Dobin et al., 2013) for alignment, or new databases can be integrated as plant-specific resources expand without altering the overall structure of the workflow. This flexibility enables seamless adaptation to new genomes, sequencing platforms, and research questions, while maintaining reproducibility and scalability for large-scale analyses.

**Fig. 1:**
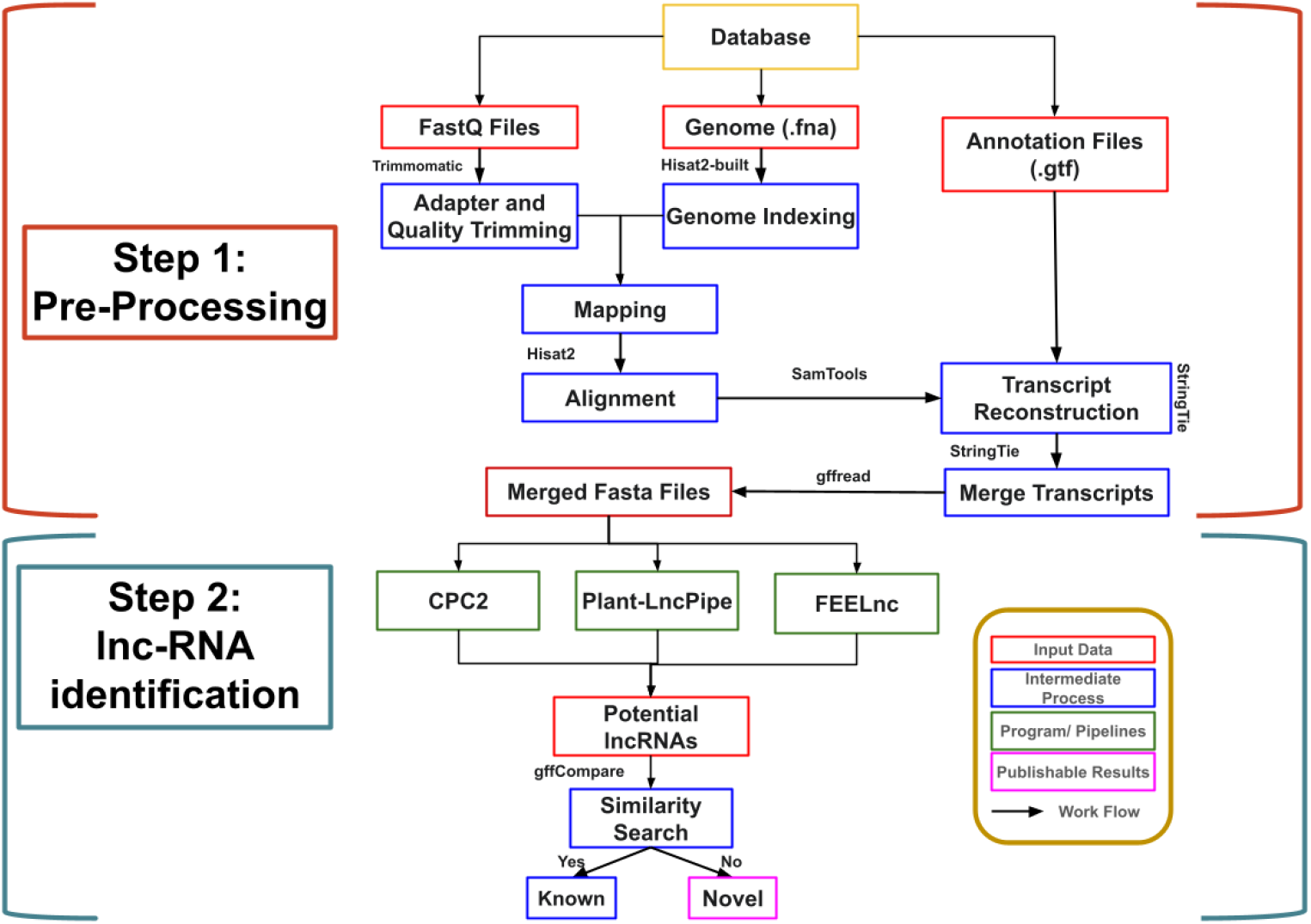
The PLncFire scalable pipeline for the identification and annotation of plant lncRNAs from transcriptomic data. Key stages include: (1) Pre-processing: quality control, adapter trimming, and read alignment; (2) lncRNA Identification: transcript assembly, filtering, and consensus coding-potential assessment using CPC2, PlantLncPipe, and FEELnc; and later classification into known/novel lncRNAs.

Performance evaluation of PLncFire was conducted by benchmarking against existing pipelines using publicly available transcriptomic datasets from NCBI (Schoch et al., 2020). Since PLncFire produces binary predictions based on a consensus of three independent coding-potential tools, the prediction quality was assessed using standard classification metrics, including accuracy, precision, recall, and F1 score, which were calculated by comparing predicted lncRNAs with reference annotations. This ensured that evaluation focused on both recovery of annotated transcripts and minimisation of false positives, providing a balanced measure of performance. By integrating sequential steps of quality control (Bolger et al., 2014; Brown et al., 2017), reference-guided transcript assembly (Shumate et al., 2022), stringent transcript filtering, consensus-based coding potential prediction (Kang et al., 2017; Wucher et al., 2017; Tian et al., 2024), and database-driven classification, PLncFire systematically progresses from raw sequencing reads to high-confidence lncRNA catalogues. This structured workflow ensures that candidate transcripts are rigorously evaluated, classified into known and novel categories, and quantified for expression profiling, providing a complete pipeline ready for downstream analysis.

## 3 Results

We benchmarked PLncFire against CPC2 (Kang et al., 2017), PlantLncPipe (Tian et al., 2024), FEELnc (Wucher et al., 2017), PlantLncBoost (Tian et al., 2025) using transcriptomic datasets from NCBI of Arachis duranensis (NCBI Accession: PRJNA258023)(Zhang et al., 2020) and Glycine max (NCBI Accession: PRJNA48389) (Chang et al., 2013). Evaluation was based on recovery of annotated lncRNAs, novel predictions, and standard statistical metrics.

lncRNA identification statistics are summarized in Figure 2, which breaks down recovered (overlapping with annotated lncRNAs), novel, other biotype, and missed lncRNAs across pipelines. In A. duranensis (Figure 2a), PLncFire predicted 10,155 lncRNAs, recovering all 8017 annotated transcripts (100%) and identifying 2138 novel candidates, without misclassification into unrelated biotypes. FEELnc predicted 8017 lncRNAs, recovering 7045 (88%) but missing 972. CPC2 recovered 3563 (44%) and PlantLncPipe only 182 (2%). By contrast, PlantLncBoost generated 28,509 predictions, recovering 6053 annotated lncRNAs (76%) but misclassifying more than 18,000 as other biotypes, reflecting substantial overprediction. A similar pattern was observed in G. max (Figure 2b), PLncFire predicted 4196 lncRNAs, recovering all 4181 annotated transcripts (100%) and reporting only 15 novel candidates. FEELnc recovered 3547 annotated lncRNAs (85%) but missed 634. CPC2 recovered 2821 (67%) and PlantLncPipe only 220 (5%). PlantLncBoost recovered just 552 annotated lncRNAs (13%) despite predicting 795 in total, with most being misassigned or spurious.

**Fig. 2:**
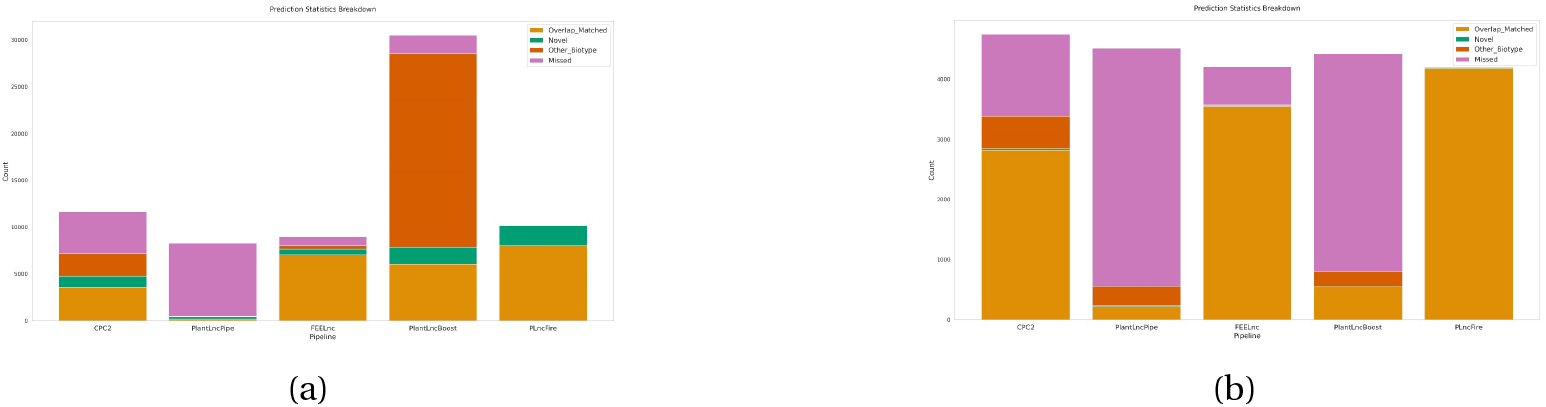
lncRNA identification statistics across pipelines. (a) Arachis duranensis and (b) Glycine max. Each embedded bar represents the distribution of recovered (overlap with annotated lncRNAs), novel candidates, transcripts classified as other biotypes, and missed lncRNAs, allowing direct comparison of pipeline performance.

Figure 3 further illustrates benchmarking accuracy in terms of recovery rates. PlncFire recovered all annotated lncRNAs in the evaluated datasets and showed higher recovery rates than the other methods considered in this study. FEELnc performed second best, but its recall remained incomplete. CPC2 and PlantLncPipe systematically under-predicted, whereas PlantLncBoost recovered a moderate fraction but suffered from instability and inflated false positives.

**Fig. 3:**
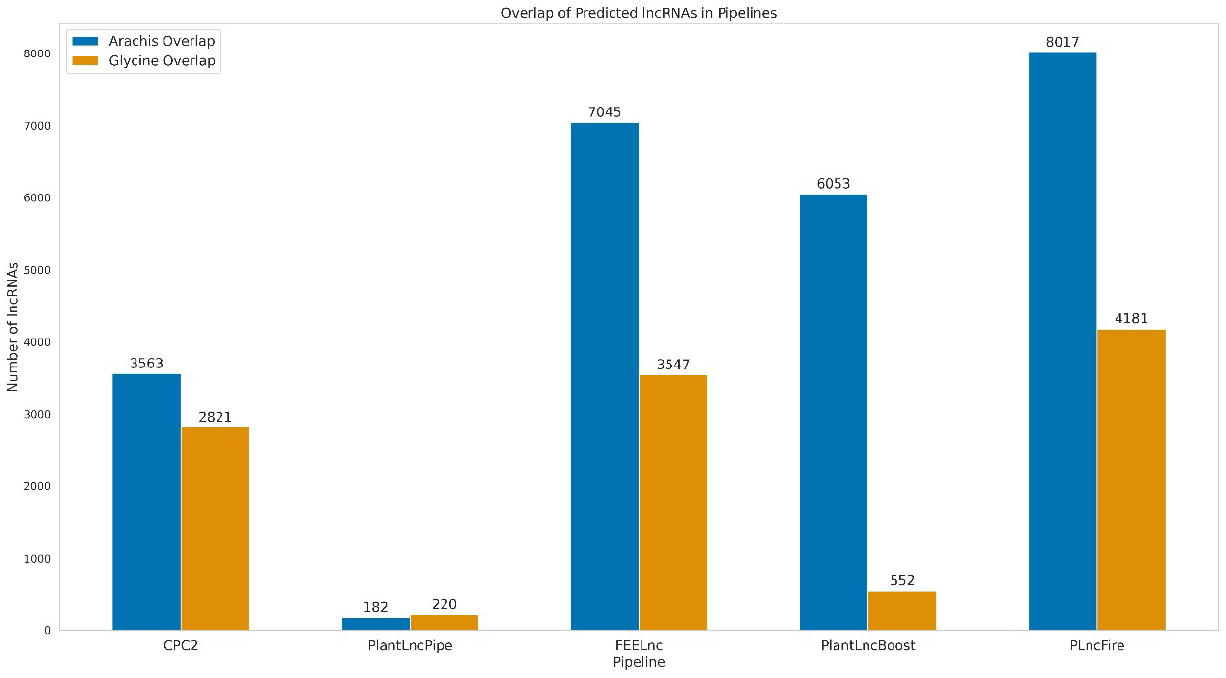
Benchmarking accuracy of lncRNA recovery in Arachis duranensis and Glycine max: comparison of PLncFire with existing pipelines.

## 4 Discussion

To capture performance in terms of classification quality, we compared pipelines across accuracy, precision, recall, and F1 score, summarized in radar charts (Figure 4a, b). In A. duranensis, PLncFire achieved perfect scores across all metrics (accuracy, precision, recall, and F1 = 1.00). FEELnc showed strong but imperfect balance (accuracy = 0.97, precision = 0.95, recall = 0.88, F1 = 0.91). CPC2 achieved accuracy of 0.84 but low recall (0.44), while PlantLncPipe performed poorly across all metrics (F1 = 0.04). PlantLncBoost displayed high recall (0.76) but very low precision (0.23), resulting in an unstable F1 score of 0.35. The same trend was evident in G. max. PLncFire achieved the highest values for the evaluated performance metrics in both datasets, followed by FEELnc (accuracy = 0.99, precision = 1.00, recall = 0.85, F1 = 0.92). CPC2 balanced moderate performance (accuracy = 0.97, precision = 0.84, recall = 0.67, F1 = 0.75), whereas PlantLncPipe showed lower performance relative to the other evaluated methods (F1 = 0.09). PlantLncBoost showed modest precision (0.70) but extremely low recall (0.13), yielding a poor F1 score of 0.22. Taken together, these benchmarks demonstrate that PLncFire consistently delivers complete recovery of annotated lncRNAs while maintaining high precision and balanced recall, a combination not achieved by any other pipeline tested. Earlier methods such as CPC2 and PlantLncPipe suffer from systematic underprediction, missing large fractions of genuine lncRNAs, whereas PlantLncBoost, despite being introduced as a machine learning–based advance, continues to display instability and excessive false positives even after recent updates. By avoiding both under and overprediction, These results suggest that PLncFire may serve as a useful framework for plant lncRNA identification and annotation. for large-scale lncRNA discovery in plants.

**Fig. 4:**
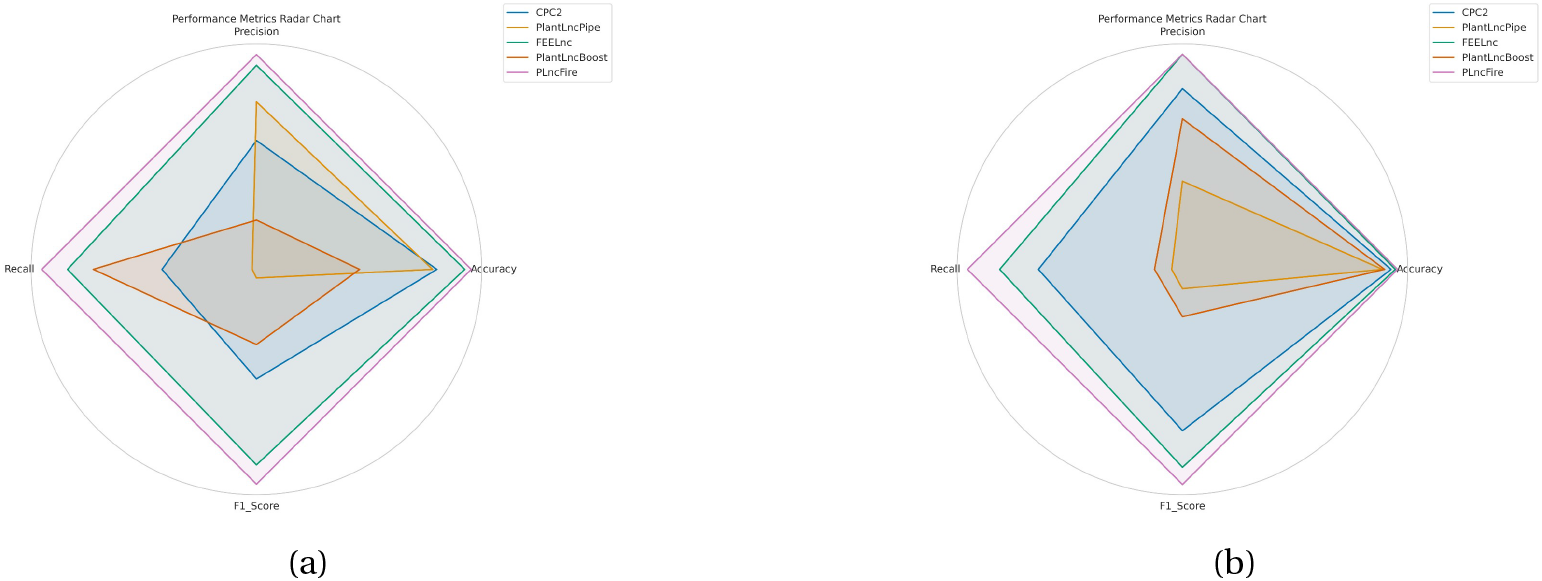
Radar charts summarizing statistical performance metrics (Recall value, Precision, F1 Score, Accuracy) for (a) Arachis duranensis and (b) Glycine max.

Although PLncFire performed well on the datasets examined in this study, some limitations remain. The evaluation was carried out using only two plant species, and therefore the results may not fully reflect the diversity of plant transcriptomes. Testing across additional species, biological conditions, and independent datasets will be necessary to better understand the broader applicability of the workflow. In addition, the present study focused on the computational identification and annotation of lncRNAs and did not include experimental validation or detailed functional characterization of the predicted candidates. Factors such as GC-content variation across plant genomes were also not specifically investigated. Consequently, the identified lncRNAs should be regarded as candidates for further biological study. Future work will focus on expanding the range of datasets evaluated and incorporating downstream analyses to further assess the utility of the pipeline.

## 5 Conclusion

Long non-coding RNAs are key regulators of plant development, stress responses, and genome evolution, yet their discovery is often limited by inconsistent methods. PLncFire provides a modular, reproducible workflow that combines stringent filtering, consensus based coding potential assessment, and comprehensive annotation. Benchmarking against PlantLncBoost demonstrates that PLncFire identified a larger set of candidate lncRNAs in the evaluated datasets while maintaining agreement with available annotations., minimizes false positives, and enables functional prioritization through classification and expression profiling. Its consensus-driven design ensures robustness, interpretability, and scalability across species and sequencing platforms. By standardizing lncRNA discovery, PLncFire offers a reliable platform for advancing plant functional genomics and uncovering regulatory complexity.

## Acknowledgment

The results presented in this paper were obtained using HPC facilities under ICE-BRAF, CDAC, Pune, India and PARAM Shavak Facility at Institute of Advanced Research (Gandhinagar, Gujarat, India) sponsored by GUJCOST.

## Declarations

### Consent to Publish

Consent to Publish: Not applicable.

### Ethics Approval and Consent to Participate

Ethics declaration: Not applicable. This study does not involve human participants, animals, or any clinical data.

### Data Availability

The datasets analyzed in this study are publicly available from the NCBI database. The source code for the PLncFire pipeline is available at https://github.com/ahsan-rizvi/PLncFire.git.

### Author Contributions

Surbhi Mistry led the study, designed and developed the PLncFire pipeline, performed all major analyses, and wrote the manuscript. Shivani Saxena contributed to pipeline development and assisted with data analysis. Dr. Ahsan Rizvi provided overall supervision, validated the pipeline and workflow, and critically reviewed and approved the manuscript. All authors read and approved the final version of the manuscript.

## References

1. Zhang, X., Wang, W., Zhu, W., Dong, J., Cheng, Y., Yin, Z., et al. (2019). Mechanisms and functions of long non-coding RNAs at multiple regulatory levels. Int J Mol Sci 20. doi: 10.3390/ijms20225573

2. Budak, H., Kaya, S. B., and Cagirici, H. B. (2020). Long Non-coding RNA in Plants in the Era of Reference Sequences. Front Plant Sci 11. doi: 10.3389/fpls.2020.00276

3. Psvv, C., Joseph, A., Ebenezer, P., Sankar, V., Suravajhala, R., Rao, R. S. P., et al. (2025). An introduction to non-coding RNAs. Prog Mol Biol Transl Sci 214, 1–17. doi: 10.1016/BS.PMBTS.2025.01.006

4. Jha, U. C., Nayyar, H., Jha, R., Khurshid, M., Zhou, M., Mantri, N., et al. (2020). Long non-coding RNAs: Emerging players regulating plant abiotic stress response and adaptation. BMC Plant Biol 20. doi: 10.1186/s12870-020-02595-x

5. Shukla, B., Gupta, S., Srivastava, G., Sharma, A., Shukla, A. K., and Shasany, A. K. (2021). lncRNADetector: a bioinformatics pipeline for long non-coding RNA identification and MAPslnc: a repository of medicinal and aromatic plant lncRNAs. RNA Biol 18, 2290–2295. doi: 10.1080/15476286.2021.1899673

6. Zhang, L., Lin, T., Zhu, G., Wu, B., Zhang, C., and Zhu, H. (2023). LncRNAs exert indispensable roles in orchestrating the interaction among diverse noncoding RNAs and enrich the regulatory network of plant growth and its adaptive environmental stress response. Hortic Res 10. doi: 10.1093/hr/uhad234

7. Chi, Y., Wang, D., Wang, J., Yu, W., and Yang, J. (2019). Long non-coding RNA in the pathogenesis of cancers. Cells 8. doi: 10.3390/cells8091015

8. Karmakar, K., Kundu, A., Rizvi, A. Z., Dubois, E., Severac, D., Czernic, P., et al. (2019). Transcriptomic analysis with the progress of symbiosis in “Crack-Entry” legume arachis hypogaea highlights its contrast with “Infection Thread” adapted legumes. Molecular Plant-Microbe Interactions 32, 271–285. doi: 10.1094/MPMI-06-18-0174-R

9. Ramakrishnaiah, Y., Kuhlmann, L., & Tyagi, S. (2020). Towards a comprehensive pipeline to identify and functionally annotate long noncoding RNA (lncRNA). Computers in Biology and Medicine, 127, 104028.

10. Mistry, S., Saxena, S., Patel, N., & Rizvi, A. (2026). A Computational Method for Predicting Tumour Cells in Arachis Hypogaea Root Nodules Through Differentially Expressed Genes. In BIO Web of Conferences (Vol. 228, p. 01004). EDP Sciences.

11. Tian, X. C., Chen, Z. Y., Nie, S., Shi T. Le, Yan, X. M., Bao, Y. T., et al. (2024). Plant-LncPipe: a computational pipeline providing significant improvement in plant lncRNA identification. Hortic Res 11. doi: 10.1093/hr/uhae041

12. Wucher, V., Legeai, F., Hédan, B., Rizk, G., Lagoutte, L., Leeb, T., … & Derrien, T. (2017). FEELnc: a tool for long non-coding RNA annotation and its application to the dog transcriptome. Nucleic acids research, 45(8), e57–e57.

13. Li, H., Handsaker, B., Wysoker, A., Fennell, T., Ruan, J., Homer, N., et al. (2009). The Sequence Alignment/Map format and SAMtools. Bioinformatics 25, 2078–2079. doi: 10.1093/bioinformatics/btp352

14. Guo, J., Gao, J., and Liu, Z. (2022). HISAT2 Parallelization Method Based on Spark Cluster., in Journal of Physics: Conference Series, (IOP Publishing Ltd). doi: 10.1088/1742-6596/2179/1/0120

15. Singh, U., Khemka, N., Rajkumar, M. S., Garg, R., & Jain, M. (2017). PLncPRO for prediction of long non-coding RNAs (lncRNAs) in plants and its application for discovery of abiotic stress-responsive lncRNAs in rice and chickpea. Nucleic acids research, 45(22), e183–e183.

16. Ritu, Gupta S., Sharma, N. K., & Shankar, R. (2021). DeepPlnc: Discovering plant lncRNAs through multimodal deep learning on sequential data. bioRxiv, 2021-12.

17. Pertea, M., and Pertea, G. (2020). GFF Utilities: GffRead and GffCompare. F1000Res 9. doi: 10.12688/f1000research.23297.1

18. Tian, X. C., Nie, S., Domingues, D., Rossi Paschoal, A., Jiang, L. B., and Mao, J. F. (2025). PlantLncBoost: key features for plant lncRNA identification and significant improvement in accuracy and generalization. New Phytologist 247, 1538–1549. doi: 10.1111/nph.70211

19. Kang, Y. J., Yang, D. C., Kong, L., Hou, M., Meng, Y. Q., Wei, L., et al. (2017). CPC2: A fast and accurate coding potential calculator based on sequence intrinsic features. Nucleic Acids Res 45, W12–W16. doi: 10.1093/nar/gkx428

20. Brown, J., Pirrung, M., and Mccue, L. A. (2017). FQC Dashboard: Integrates FastQC results into a web-based, interactive, and extensible FASTQ quality control tool. Bioinformatics 33, 3137–3139. doi: 10.1093/bioinformatics/btx373

21. Danecek, P., Bonfield, J. K., Liddle, J., Marshall, J., Ohan, V., Pollard, M. O., et al. (2021). Twelve years of SAMtools and BCFtools. Gigascience 10. doi: 10.1093/gigascience/-giab008

22. Shumate, A., Wong, B., Pertea, G., and Pertea, M. (2022). Improved transcriptome assembly using a hybrid of long and short reads with StringTie. PLoS Comput Biol 18. doi: 10.1371/journal.pcbi.1009730

23. Sweeney, B. A., Petrov, A. I., Burkov, B., Finn, R. D., Bateman, A., Szymanski, M., et al. (2019). RNAcentral: A hub of information for non-coding RNA sequences. Nucleic Acids Res 47, D221–D229. doi: 10.1093/nar/gky1034

24. Szcześniak, M. W., and Wanowska, E. (2024). CANTATAdb 3.0: An Updated Repository of Plant Long Non-Coding RNAs. Plant Cell Physiol 65, 1486–1493. doi: 10.1093/pcp/pcae081

25. Jin, J., Lu, P., Xu, Y., Li, Z., Yu, S., Liu, J., et al. (2021). PLncDB V2.0: A comprehensive encyclopedia of plant long noncoding RNAs. Nucleic Acids Res 49, D1489–D1495. doi: 10.1093/nar/gkaa910

26. Camacho, C., Coulouris, G., Avagyan, V., Ma, N., Papadopoulos, J., Bealer, K., et al. (2009). BLAST+: Architecture and applications. BMC Bioinformatics 10. doi: 10.1186/1471-2105-10-421

27. Dobin, A., Davis, C. A., Schlesinger, F., Drenkow, J., Zaleski, C., Jha, S., et al. (2013). STAR: Ultrafast universal RNA-seq aligner. Bioinformatics 29, 15–21. doi: 10.1093/bioinformatics/bts635

28. Schoch, C. L., Ciufo, S., Domrachev, M., Hotton, C. L., Kannan, S., Khovanskaya, R., et al. (2020). NCBI Taxonomy: A comprehensive update on curation, resources and tools. Database 2020. doi: 10.1093/database/baaa062

29. Bolger, A. M., Lohse, M., and Usadel, B. (2014). Trimmomatic: A flexible trimmer for Illumina sequence data. Bioinformatics 30, 2114–2120. doi: 10.1093/bioinformatics/btu170

30. Chang, S., Wang, Y., Lu, J., Gai, J., Li, J., Chu, P., et al. (2013). The Mitochondrial Genome of Soybean Reveals Complex Genome Structures and Gene Evolution at Intercellular and Phylogenetic Levels. PLoS One 8. doi: 10.1371/journal.pone.0056502

31. Zhang, R., Wang, Y. H., Jin, J. J., Stull, G. W., Bruneau, A., Cardoso, D., et al. (2020). Exploration of Plastid Phylogenomic Conflict Yields New Insights into the Deep Relationships of Leguminosae. Syst Biol 69, 613–622. doi: 10.1093/sysbio/syaa013

